# Genomic Analysis of Powdery Mildew Resistance in a Hop (*Humulus lupulus* L.) Bi-parental Population Segregating for “R6-locus”

**DOI:** 10.1101/864249

**Authors:** Lillian K. Padgitt-Cobb, Sarah B. Kingan, John A. Henning

## Abstract

Genetic response in hop to fungal pathogen infection has been evaluated at the chromosomal level through QTL analyses but very little information exists on the expression of genes during infection periods. Raw GBS reads and phenotypic data from a previously published QTL analysis along with a newly assembled PacBio-derived hop reference genome were used to re-evaluate resistance to races v4/v6 of powdery mildew (PM; *Podosphaera humuli*). QTL analyses revealed two tightly linked regions of association on a single linkage group. The three SNP markers most tightly linked to PM resistance (found on contig 000559F) were observed downstream from a putative R-gene locus for powdery mildew resistance. This 230 kb region contained a series of seven putative R-genes surrounded by seven putative peroxidase-3 genes downstream and seven putative glucan endo-1,3-beta-glucosidase upstream and an expressed F-box domain protein. RNAseq data showed all putative R-genes along with all putative glucan endo-1,3-beta-glucosidase genes were expressed under diseased conditions, while none of the peroxidase genes were expressed. The second region contained three SNPs found on contig 002916F next to two putative R-genes. RNAseq data showed complex expression of exons contained in putative isoforms of R-genes. This preliminary information will prove valuable information for development of precise markers located either within or next to genes responsible for race v4/v6 PM resistance.

## INTRODUCTION

Hop (*Humulus lupulus* L var lupulus) is a diploid (2n=2x=18 + XX/XY) dioecious perennial crop species used principally for beer bittering and flavoring although alternate uses include replacement of antibiotics in poultry (Bortoluzzi et al., 2014), anti-cancer (Miranda et al., 1999, Jiang et al., 2018), metabolic disorder (Miranda et al., 2016) and estrogen replacement therapy (Hemachandra, et al., 2012). It is produced on six meters tall trellis under wide row spacing (three to five meters) between rows in most countries with the exception of Great Britain where some hops are produced under 2.5 m low-trellis using hops bred specifically for these conditions (Neve, 1991). The harvested portion of the hop plant is the mature female hop flower or “cone” while male plants are used only for breeding purposes.

The greatest constraints for hop production are water, fertility and disease management. Timing and availability of water and fertilizer applications are easily controlled factors that growers have addressed with implementation of drip irrigation systems in the Pacific Northwest region of USA. However, disease management is not a trivial factor in hop production as many of the most popular hop cultivars are susceptible to the two most prevalent diseases: hop downy mildew (caused by: *Pseudoperonospora humuli* (Miyabe & Takah.) G.W. Wilson) and hop powdery mildew (caused by: *Podosphaera macularis* (Wallr.) U. Braun & S. Takam). Pathogen resistance to chemical treatments has been documented (Klein, 1994; Gent et al., 2008) and efforts towards implementation of physical control methods such as springtime pruning are limited due to the potential spread of virus and viroid’s using mechanical pruning. With these factors in mind, it follows that the most economical means of controlling fungal diseases is to produce hop cultivars that are resistant to fungal pathogens.

Strategies for plant resistance breeding have focused traditionally upon the identification of, and selection for, complete or qualitative resistance—usually expressed as a single dominant resistance-gene or “R-gene” (Flor, 1971), which is defined as a “gene-for-gene” resistance whereby the host plant possessed a single gene that conferred resistance to a single virulence gene (avr) found within the pathogen. In almost all cases, R-genes were race specific within a single pathogen species and did not confer broad-scope resistance to other fungal or bacterial pathogens. Studies within the past two decades have shown R-gene response to infection as a more complex system than a single gene responsible for resistance and several pathways for resistance, as well as multi-pathogen resistance, has been described (Feys and Parker, 2000; Dangle & Jones, 2001). R-gene function and activity were recently evaluated and summarized by Kourelis and van der Hoom (2017).

Powdery mildew (PM) resistance in several plant species has been shown in many cases to be under qualitative resistance with well-defined R-genes (Donald et al. 2002; Consonni et al., 2006). Resistance to powdery mildews in barley has also been attributed to loss of function in *mlo* genes (mildew resistance locus O). This latter means of resistance has been speculated to be more durable than resistance based solely on the presence of a single R-gene (Consonni et al., 2006)

Previous research on hop powdery mildew (PM) suggests that resistance is based primarily upon qualitative expression with seven known R-genes sources (Neve, 1991; Henning et al., 2017; Wolfenberger et al., 2016). QTL analyses on hop powdery mildew resistance performed over the past decade have revealed several possible locations for R-genes (Henning et al., 2011; Henning et al., 2017; McAdam et al, 2013; Cerenak, 2006) although no genes with homology to known “R-genes” were identified. Unfortunately, different marker systems were used across all these studies and there are no means of comparing results although Henning et al. (2017) used genome-based SNP markers and future studies using their genomic data can be compared for genomic locations of R-genes. An important finding in most of these studies showed the presence of highly significant QTL but also showed the presence of QTLs having smaller impact upon expression of disease resistance, leading to the idea that resistance to PM may be conditioned by both qualitative and quantitative genic control (Henning, et al. 2017; Wolfenberger et al., 2016).

Henning et al. (2017) based their identification of SNPs linked to PM resistance upon the reference ‘Teamaker’ genome (Hill et al. 2016) and the resulting genetic map was used for QTL identification. The Teamaker genome only covers 1.7 Gb of the projected haploid size of 2.8 Gb (Hill et al., 2016). Thus, a large portion of the genome that was used for SNP identification was missing and the resulting genetic map developed for QTL analysis could potentially miss important gene-space regions. The gene-space both within and surrounding QTLs in Henning et al (2017) were evaluated for genes that were potentially involved in resistance to powdery mildew. No R-genes were identified although two genes involved in the biosynthesis of chalcone synthase were putatively identified. Chalcone synthase is an important pre-cursor for the flavonoid/isoflavonoid biochemical pathway. It is known to be involved in responses to biotic and abiotic stress and in particular is involved in the salicylic acid (SA) defense pathway (Dao et al., 2011). However, it remains that genes directly involved in PM resistance have not been identified in hop.

Genomic evaluation of plant resistance to PM in hop is needed as an aid towards developing breeding strategies to these important fungal diseases. This preliminary study was designed to begin elucidating the genes potentially involved in resistance to a specific race of PM.

## MATERIALS AND METHODS

### Plant Material

One bi-parental mating population was developed as previously reported (Henning et al., 2017). The population, USDA 2000001, was derived from a cross between the PM-resistant (R-4, R-6) USDA cultivar ‘Newport’ (Henning et al. 2004) and PM-susceptible male germplasm USDA 21110M. Seeds were treated with cold (~5°C), moist conditions for eight weeks to break dormancy. Seed planting and germination took place in a heated glasshouse (~25°C day and night) with 16 hours under light and eight hours dark. A sulfur-emitter was used constantly within the glasshouse to prevent any disease until inoculation. Replicates of plants were developed using soft-wood cuttings planted in Oasis planting trays (https://www.oasisgrowersolutions.com/ verified 10/01/2019).

### Disease Inoculations and Scoring

All plant materials were inoculated with known quantitates of PM as described in Henning et al. (2017). Seventy-five offspring and parents were grown in a replicated block design (three blocks in time) with two clones per genotype in a glasshouse under controlled environmental conditions designed to maximize expression of disease symptoms of susceptible offspring and parents. Disease scores for each genotype (two clones per genotype) within a block were averaged. Analysis of variance was performed with blocks and genotypes as factors. Significant differences among genotypes were observed but there were no significant differences between blocks. As a result, disease scores for genotypes were averaged across blocks.

Scoring for PM resistance levels under glasshouse conditions were previously described in Henning et al. (2017). In summary, plants were scored with a six-step ordinal scale (0-5) where 0 = no disease symptoms (highly resistant); 1 = necrotic flecks, non-sporulating blisters, or aborted infection (resistant); 2 = one or few small lesions on plant with only slight sporulation (moderately resistant); 3 = multiple lesions on a plant, but not on all susceptible leaves (moderately susceptible); 4 = multiple lesions on all susceptible leaves (susceptible); 5 = coalescing lesions on multiple leaves (highly susceptible).

### Genotyping-by-sequencing, QTL Analysis and Association Studies

All plant materials (parents and offspring) for DNA samples were grown under glasshouse conditions with all attempts to control pathogen and insect infestation as described by Henning et al. (2017). A modified extraction protocol using Qiagen DNAeasy Mini extraction kits (QIagen Incorporated, Germantown, MD) was used to extract DNA from young hop leaves as described by Henning et al. (2016). Genotyping-by-sequencing was performed as described by Elshire et al (2011) using an Illumina HiSeq 3000 sequencing. SNP identification was performed as described by Henning et al. (2017) with the exception that a PacBio (PacBio Sequel Sequencing Instrument; Pacific Biosciences, Menlo Park, CA) sequenced and Falcon-assembled (Chin et al., 2016) hop genome—based upon the hop cultivar ‘Cascade’ (http://hopbase.cgrb.oregonstate.edu)—was used.

The genetic map for QTL studies was developed from SNP markers identified via TASSEL v5.0 pipeline (Elshire et al. 2011) using the PacBio ‘Cascade’ genome as reference and the raw GBS sequencing data from Henning et al. (2017). The resulting markers for this map were pre-filtered in TASSEL version 5.2.43 (http://www.maizegenetics.net/tassel; verified 05/02/2018) for a genetic depth (2X or greater) and presence in 90% of genotypes and in Microsoft Excel for Mac v16.16.4 (Microsoft Inc; Redmond, WA) for fit with expected genetic segregation: 1:1 for AA x Aa and 1:2:1 for Aa x Aa. JoinMap Version 5.0 (64-bit; www.kyazma.nl) was used for linkage group assignment and mapping of markers using the same procedures as previously reported (Henning et al., 2017) with modifications for determining the final map used for QTL analysis. Both “Regression Mapping” (REG) as well as “Maximum Likelihood” (ML) procedures were used to estimate genetic maps. With REG mapping, only two rounds of estimation for marker location and distance were utilized rather than three and reasonable distances for overall length of linkage groups were obtained. Maximum likelihood was then utilized on all linkage groups using default settings in JoinMap 5.0. The overall length of resulting linkage groups from ML were elevated over that obtained for REG mapping. However, grouping of SNP markers located on the same contig (the actual physical map) using ML mapping was superior over that obtained from REG mapping with REG mapping typically breaking up physically-linked SNPs into different regions of the linkage group. Because of these two observations, the distances between markers and overall length of linkage groups was transformed using the following calculation:

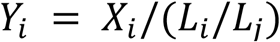

With *Y_i_* the adjusted location of marker *i, X_i_* the location of marker *i* using ML mapping, *L_i_* the overall length of linkage group containing marker *i* using ML mapping, and *L_i_* the overall length of linkage group containing marker *i* using REG mapping. This transformation aligned all markers across all linkage groups into distances/locations that were relevant to the overall length of the linkage groups observed using REG mapping. The transformed mapping distances (ie: using REG linkage group distances to modify ML linkage group distances) were used for all subsequent QTL evaluations.

The final map used for QTL analyses consisted of 2045 markers across 10 linkage groups (9 autosomes and 1 pseudo-autosomal group). The program, “Rqtl” (http://www.rqtl.org/; verified 8/12/18) was utilized to run interval mapping (IM; Lander and Botstein, 1989). WinQTL ver 2.5_11 (Silva et al., 2012; https://brcwebportal.cos.ncsu.edu/qtlcart/WQTLCart.htm; verified 8/12/2018) was used to verify IM as well as run composite interval mapping (CIM: Jansen and Stam, 1994). Significant threshold LOD (logarithm of the odds) values (significant > 3.2) for both IM and CIM were determined by permutations set to 500 with default values for all other settings.

### Gene Prediction on Contigs Found Within QTLs

We utilized Geneious Pro 11.0.5 (https://www.geneious.com) to identify putative genes involved in disease resistance. Contigs containing SNPs that were observed within a QTL were annotated with RNAseq data from Cascade stem, leaf and apical meristem (Padgitt-Cobb et al. 2019). Gene prediction was performed on all contigs containing significant SNPs using the Augustus (Stanke and Morgenstern. 2005) as an internal plugin. We used models trained for Arabidopsis for gene prediction. The UniProt database for “Embryophyta” (https://www.uniprot.org/taxonomy/3193) was downloaded locally and set up internally within Geneious Pro as a genomic resource from which to run BLAST searches (Basic Local Alignment Search Tool—Altshul et al. 1990) within Geneious Pro. Settings for BLAST were default settings. DNA regions, where Augustus predicted the presence of genes, were submitted for local searches by BLASTx using the UniProt Embryophyta database to determine potential homology with known genes. We used E-values of 1e-50 as threshold criteria for putative gene calls. RNAseq annotations were used as additional evidence for gene prediction in cases where E-values for Augustus-called putative genes had homologies with known genes that did not meet the 1e-50 threshold but were expressed in hop tissue. DNA regions covered by RNAseq alignments were also submitted for BLASTx local searches and following the same threshold criteria used as additional support for gene calls.

## RESULTS

### Phenotype

As previously reported (Henning et al., 2017) the phenotypic data across the two experimental replications were not significantly different from one another and as a result, data were averaged across replications for analyses. Blocks within replications were maintained for IM and CIM analyses in QTL studies and for association studies. Figure 1 shows the distribution of values for PM scores. A near-segregation of 1:1 is observed if scores of 0 (no disease) to 0.5 are considered “resistant” while values of 1-5 are susceptible.

**Figure 1.**
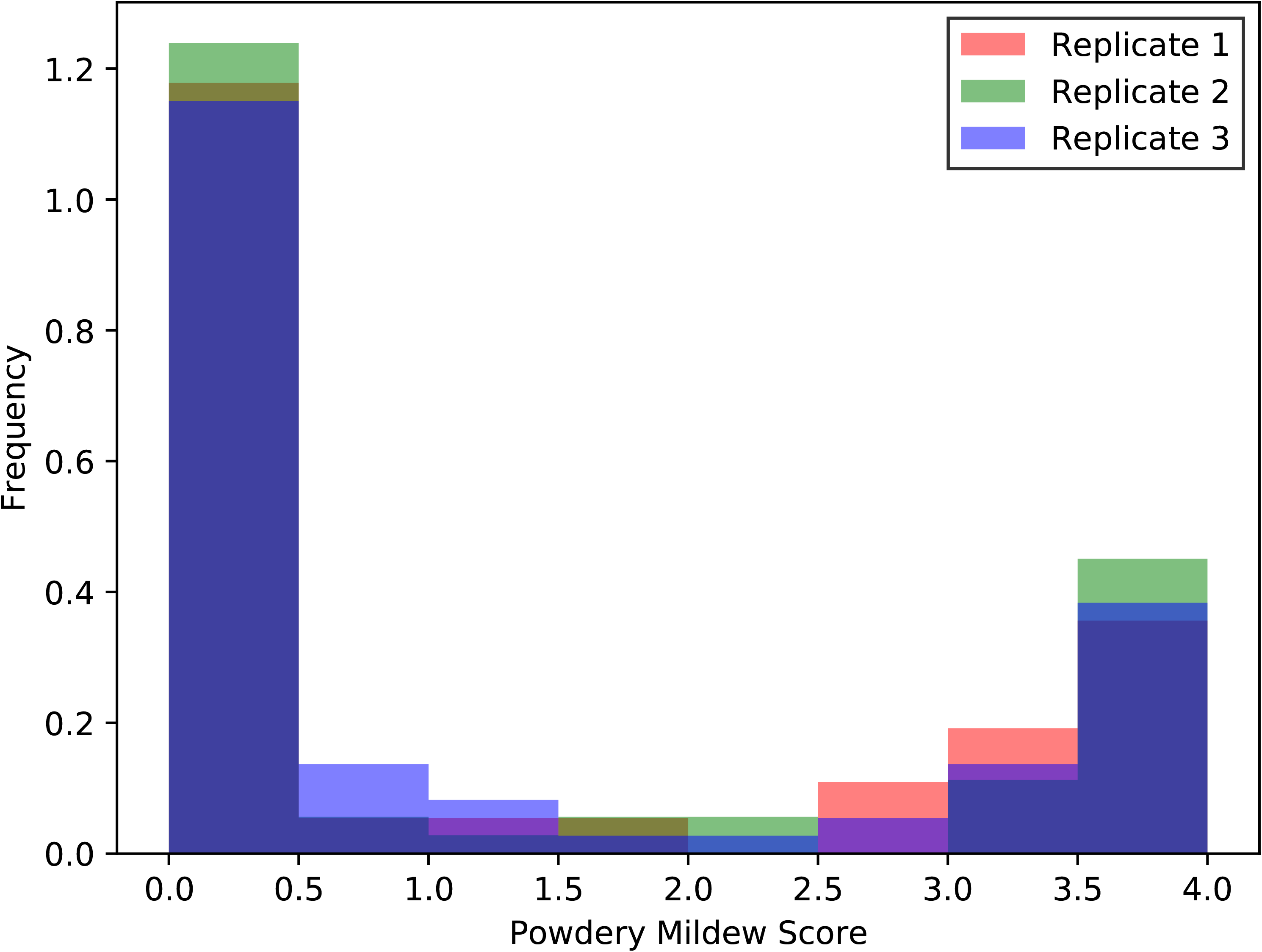
Histogram of phenotype distribution of powdery mildew response across a bi-parental mapping population between USDA Newport x USDA 21110M.

### SNP Data Set

Use of TASSEL v3.0 SNP pipeline—using the Cascade PacBio genome as reference— resulted in a data set consisting of nearly 950K filtered markers. Further filtration using TASSEL v 5.2.43 for marker depth (2x or greater) as well as markers being present in 90% of all individuals (including parents) resulted in a working data set of approximately 60K SNP markers with representative markers from almost all contigs present in the original data set (PacBio Cascade assembly statistics: 11,705 contigs with smallest contig = 20 kb, longest contig = 8.2 Mb, NG50 = 0.867 Mb http://hopbase.cgrb.oregonstate.edu/downloadDraftCascade.php).

The 60K SNP data set was imported into Microsoft Excel and markers tested for segregation fit as either testcross (aa x Aa) using Newport as test parent or the testcross (Aa x aa) with 21110M as test parent. In addition, we tested markers for segregation fit with F2-segregation (Aa x Aa). Only markers having chi-square tests greater than p = 0.05 were chosen for inclusion in map development. These tests selected approximately 8K markers that were then used for development of the genetic map (~2000 Aa x Aa markers; ~2800 Aa x aa markers; ~3100 aa x Aa markers).

### Genetic Map

The genetic map developed for this population consisted of 2045 SNP markers across 10 linkage groups (Figure 2A; Supplementary Data Tables A&B). The total distance using ML mapped spanned 21060 cM with the longest linkage group measuring 6908 cM (linkage group 10) and the shortest 850 cM for linkage group 9. Average distance between markers using ML mapping was 10.3 cM. Genetic distances of linkage groups using REG mapping spanned a total of 676.5 cM with the longest linkage group spanning 120.5 cM (linkage group 8) and the shortest spanning 34.2 cM (linkage group 7). Average distance between markers using REG mapping was 0.32 cM.

**Figure 2.**
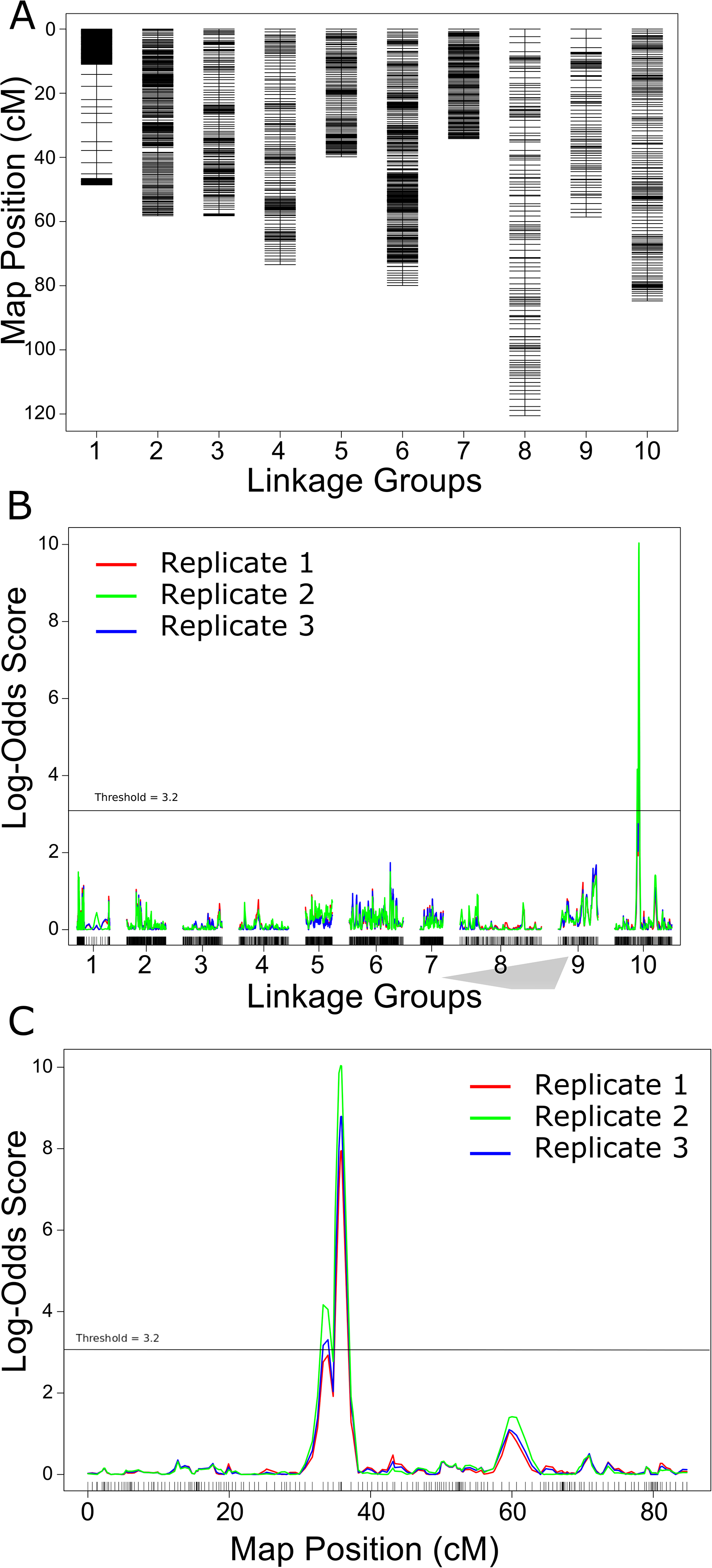
Linkage mapping and composite interval mapping results. A) Linkage groups and size across ten linkage groups (9 autosomes + 1 pseudo-autosomal region). B) Composite interval mapping (CIM) of three replicate experiments looking at the phenotype for PM resistance response for the bi-parental mapping population USDA Newport x USDA 21110M. C) Zoom in on linkage group 10 showing LOD response across three replicates in composite interval mapping.

The genetic map developed using ML mapping showed better contiguity of markers found on the same contigs as compared to the genetic map developed using REG mapping. In the case of REG mapping, many of the contiguous markers on the same contig were separated by markers present on different contigs in the linkage map. Thus, ML mapping performed better at establishing marker order along linkage groups while REG mapping performed better at estimating genetic distance between markers.

### QTL Analysis

Single marker QTL analyses identified eight markers with R^2 values greater than 0.10 with six of these having R^2 values greater than 0.20 and p-values less than 0.01 (Table 1). These six markers were observed within a span of 33.29 – 35.89 cM on linkage group 10. Four additional markers were identified having p-values ≤ 0.01 and R^2 ≥ 0.10 and were observed on linkage groups 4 & 9 (Table 1). CIM analyses (Figure 2B) eliminated markers identified in single marker QTL analyses on linkage groups 4 and 9 and focused association with PM resistance upon a single QTL located at 33.29 – 35.89 cM on linkage group 10. The first contig located within this region (contig 002196F) includes four markers located at nucleotide positions 326400 – 522644 (Figure 2C) and the second contig (contig 000559F) has three SNP markers located in nucleotide positions 178140 – 410159 (Figure 2C). No significant QTLs were observed on other linkage groups in CIM analysis.

**Table 1.**
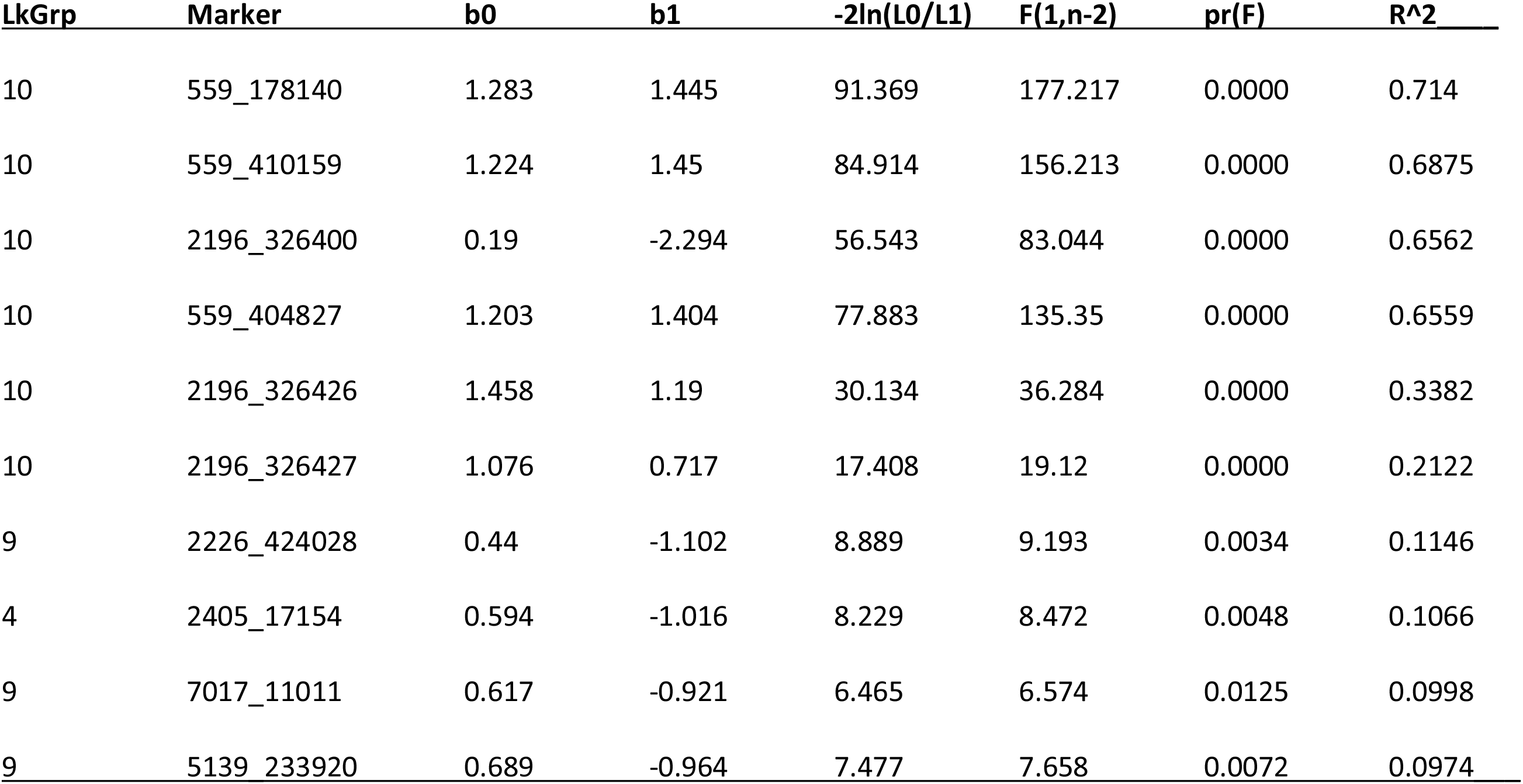
Single marker QTL analysis of PM resistance with analyses fitting the data to the simple linear regression model; y = b0 + b1*X + e. F-statistic has 1, n-2 degrees of freedom.

### Gene Identification

Contig 002196F spans a length of 567 kbp and contains 19 putative genes (Table 2) identified by Augustus and local Blastx searches against UniProt Embryophyta gene data set (https://www.uniprot.org/taxonomy/3193). Three of these putative genes have homology to R-genes found in other plant species although only two of these appear to be complete (002196F: 39443-41061 and 002196F: 46739-54401) (Figure 3). The third putative R-gene contains a retrovirus pol protein insertion and is masked on the HopBase.org Cascade genome assembly.

**Figure 3.**
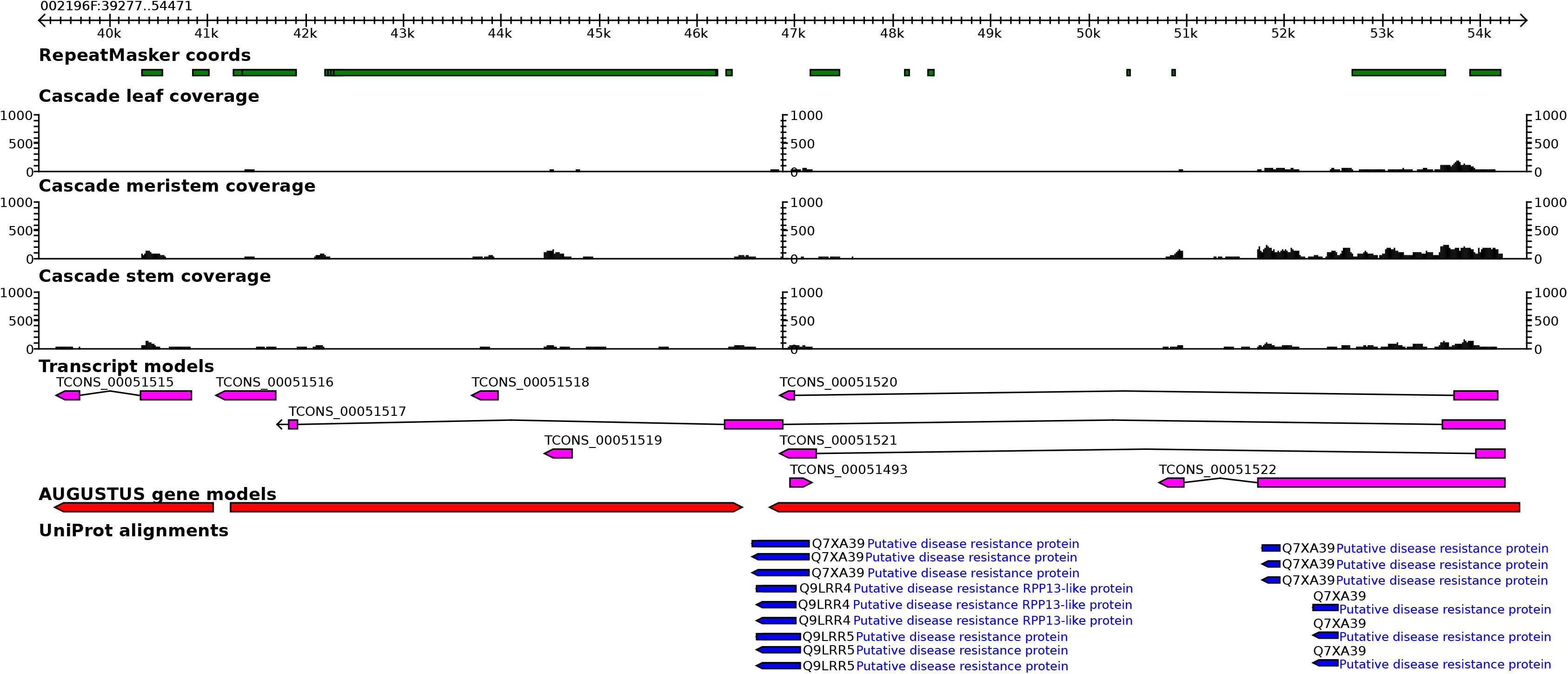
Extracted region of contig 002196F covering putative R-genes linked to expression of resistance to PM as shown in the Cascade draft genome on Hobpase.org (http://hopbase.cgrb.oregonstate.edu). Red color alignments represent Augustus-predicted genes while pink represents mRNA transcripts and blue represents Uniprot (https://www.uniprot.org/) alignments. Quantitative expression levels (black color graphs) were all normalized between tissues.

**Table 2.**
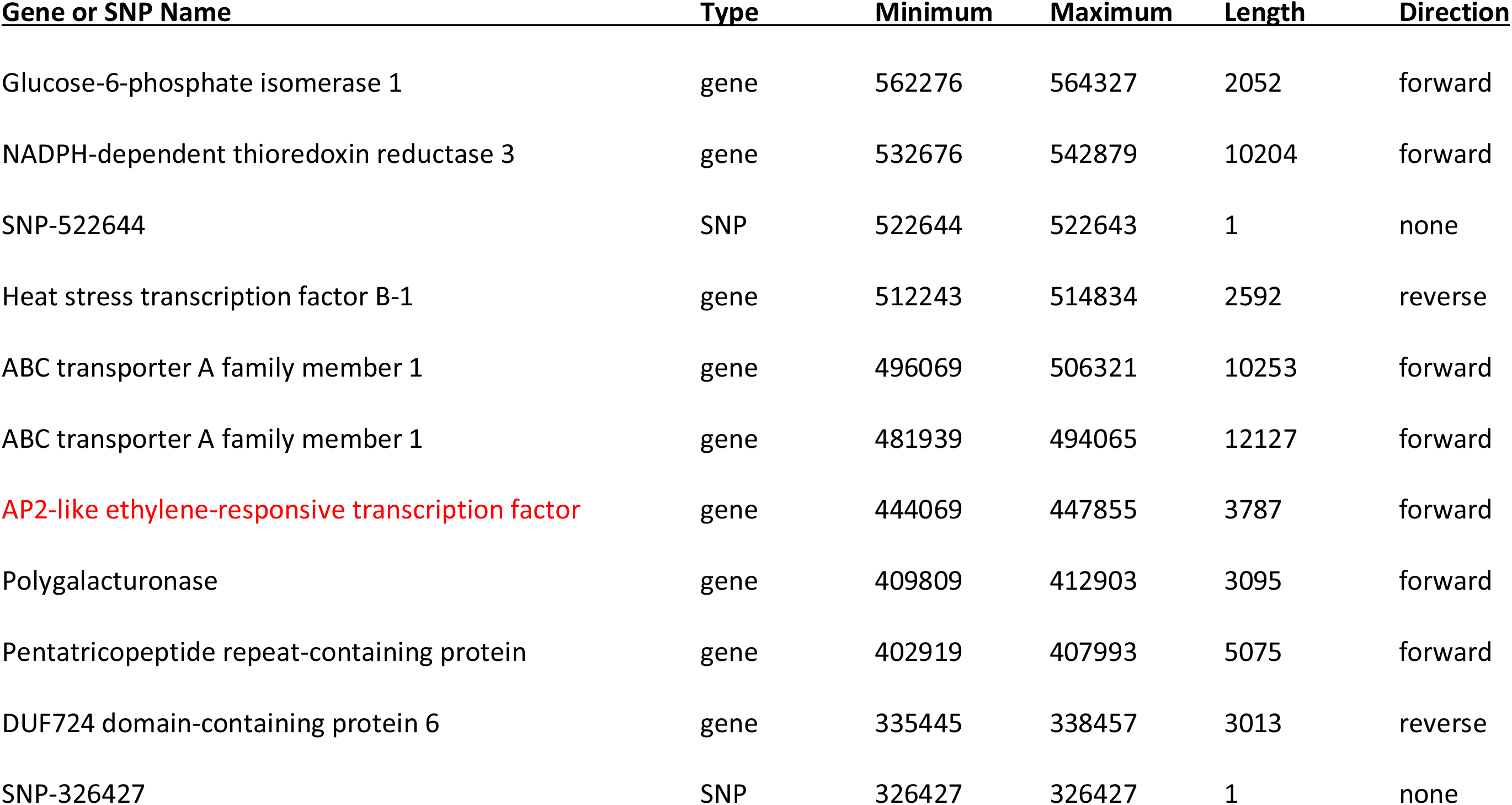
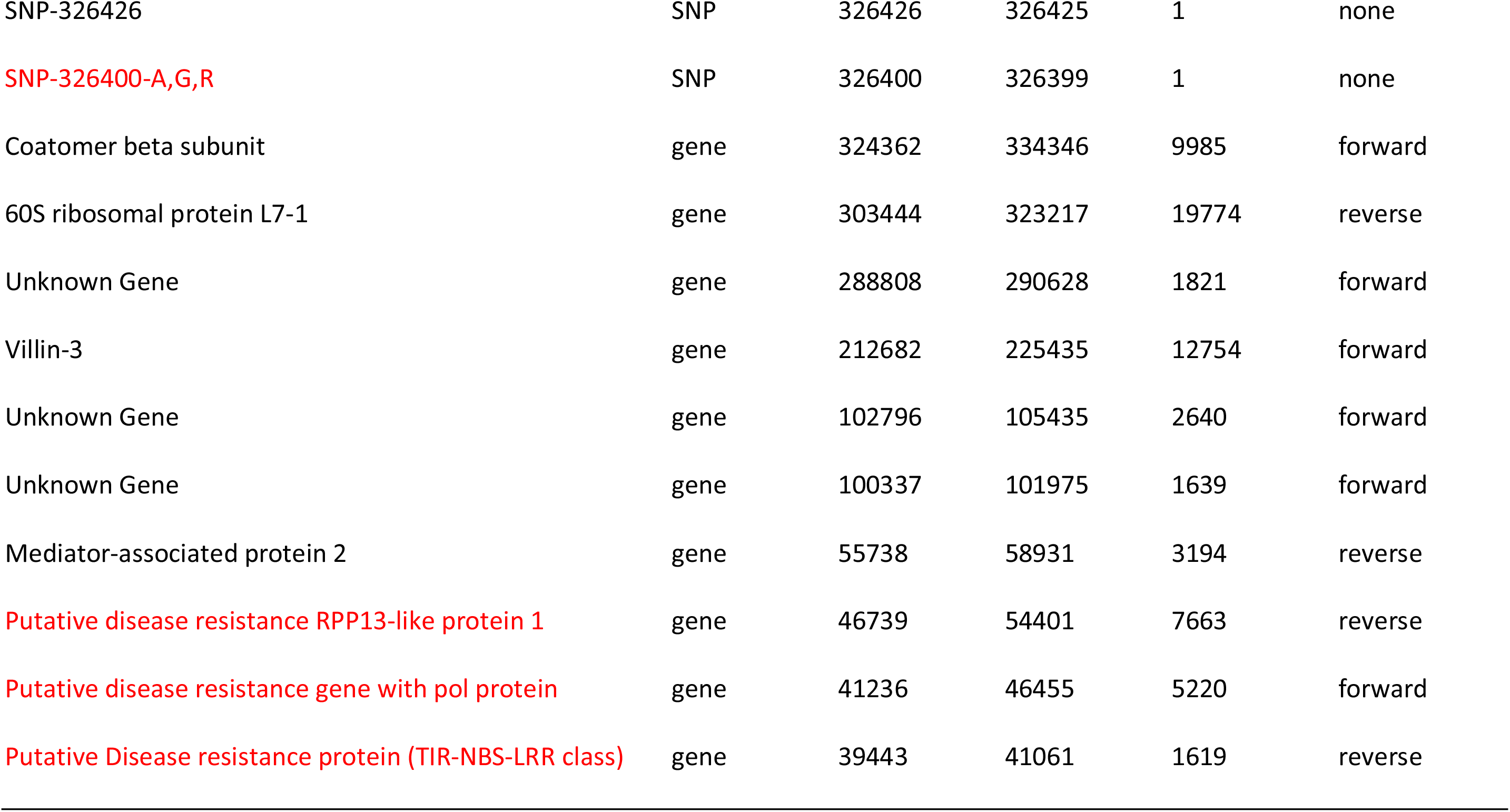
List of genes and significant SNPs found on contig 002196F. Genes with red font are expressed pathogenesis-related genes while the SNP with red font had highest significant association with PM resistance. Letters following SNP-326400 are the genotypes possible for that SNP.

| Gene or SNP Name | Type | Minimum | Maximum | Length | Direction |
| --- | --- | --- | --- | --- | --- |
| Glucose-6-phosphate isomerase 1 | gene | 562276 | 564327 | 2052 | forward |
| NADPH-dependent thioredoxin reductase 3 | gene | 532676 | 542879 | 10204 | forward |
| SNP-522644 | SNP | 522644 | 522643 | 1 | none |
| Heat stress transcription factor B-1 | gene | 512243 | 514834 | 2592 | reverse |
| ABC transporter A family member 1 | gene | 496069 | 506321 | 10253 | forward |
| ABC transporter A family member 1 | gene | 481939 | 494065 | 12127 | forward |
| AP2-like ethylene-responsive transcription factor | gene | 444069 | 447855 | 3787 | forward |
| Polygalacturonase | gene | 409809 | 412903 | 3095 | forward |
| Pentatricopeptide repeat-containing protein | gene | 402919 | 407993 | 5075 | forward |
| DUF724 domain-containing protein 6 | gene | 335445 | 338457 | 3013 | reverse |
| SNP-326427 | SNP | 326427 | 326427 | 1 | none |
| SNP-326426 | SNP | 326426 | 326425 | 1 | none |
| SNP-326400-A,G,R | SNP | 326400 | 326399 | 1 | none |
| Coatomer beta subunit | gene | 324362 | 334346 | 9985 | forward |
| 60S ribosomal protein L7-1 | gene | 303444 | 323217 | 19774 | reverse |
| Unknown Gene | gene | 288808 | 290628 | 1821 | forward |
| Villin-3 | gene | 212682 | 225435 | 12754 | forward |
| Unknown Gene | gene | 102796 | 105435 | 2640 | forward |
| Unknown Gene | gene | 100337 | 101975 | 1639 | forward |
| Mediator-associated protein 2 | gene | 55738 | 58931 | 3194 | reverse |
| Putative disease resistance RPP13-like protein 1 | gene | 46739 | 54401 | 7663 | reverse |
| Putative disease resistance gene with pol protein | gene | 41236 | 46455 | 5220 | forward |
| Putative Disease resistance protein (TIR-NBS-LRR class) | gene | 39443 | 41061 | 1619 | reverse |

The second contig observed within the QTL on linkage group 10 was found at 34.7 – 36.9 cM. Contig 000559F covers 1113 kbp and contains 50 putative genes (Table 3) as determined by Augustus and local Blastx searches. The interesting observation for this QTL is the organization of this genomic region with seven putative R-genes surrounded by an upstream grouping of putative glucan endo-1,3-beta-glucosidase genes and downstream by a set of putative peroxidase-27 genes.

**Table 3.** List of genes and SNPs found on contig 000559F. Putative genes were initially identified using Augustus and then the sequences found within the putative genes were assayed through BLAST to find homologous genes from other species that matched the hop sequence. Putative genes that are homologous to disease resistance genes in other species are printed in red font.

| **Gene or SNP Name** | **Type** | **Start** | **Finish** | **Length** | **Direction** |
| --- | --- | --- | --- | --- | --- |
| Gibberellin-regulated protein 13 | gene | 31734 | 32611 | 878 | reverse |
| Pentatricopeptide repeat-containing protein At4g32450 | gene | 42397 | 45331 | 2935 | reverse |
| N-acetylglucosaminyl transferase component | gene | 48589 | 54161 | 5573 | reverse |
| Cytochrome P450 98A2 | gene | 59456 | 61231 | 1776 | forward |
| GTP-binding protein Brassinazole insensitive pale green 2 | gene | 80629 | 84431 | 3803 | reverse |
| Syntaxin-81 | gene | 86416 | 89601 | 3186 | reverse |
| WPP domain-interacting tail-anchored protein 1 | gene | 89876 | 90528 | 653 | forward |
| WPP domain-interacting tail-anchored protein 1 | gene | 90642 | 90971 | 330 | reverse |
| Glycosyltransferase family protein 64 C3 At1g80290 | gene | 91044 | 92151 | 1108 | reverse |
| ABC transporter C family member 3 | gene | 97574 | 104451 | 6878 | reverse |
| ABC transporter C family member 3 | gene | 106116 | 112830 | 6715 | forward |
| Peroxidase 27 ¶ | gene | 154036 | 156681 | 2646 | reverse |
| Peroxidase 27 ¶ | gene | 167156 | 170621 | 3466 | reverse |
| Peroxidase 27 ¶ | gene | 176322 | 178869 | 2548 | reverse |
| SNP-178140 | SNP | 178140 | 178139 | 1 | none |
| Peroxidase 27 ¶ | gene | 218063 | 220609 | 2547 | reverse |
| Peroxidase 27 ¶ | gene | 268869 | 272689 | 3821 | reverse |
| Casein kinase 1-like protein HD16 § | gene | 306912 | 309979 | 3068 | reverse |
| Peroxidase 27 ¶ | gene | 327702 | 330219 | 2518 | reverse |
| Peroxidase 27 ¶ | gene | 379882 | 386688 | 6807 | reverse |
| Probable F-box protein At5g04010 ¶ | gene | 404293 | 405295 | 1003 | forward |
| Histone acetyltransferase type B, catalytic subunit | gene | 406873 | 412192 | 5320 | forward |
| SNP-410159 | SNP | 410159 | 410158 | 1 | none |
| Peptide chain release factor PrfB3 | gene | 432093 | 435859 | 3767 | forward |
| Folylpolyglutamate synthetase | gene | 474810 | 490099 | 15290 | forward |
| Putative disease resistance protein RGA4 § | gene | 491369 | 497348 | 5980 | reverse |
| Putative disease resistance RPP13-like protein 1 § | gene | 498943 | 502378 | 3436 | reverse |
| Putative disease resistance protein RGA1 § | gene | 518699 | 522568 | 3870 | reverse |
| Disease resistance protein RGA2 § | gene | 539261 | 543789 | 4529 | reverse |
| Putative disease resistance protein RGA1 § | gene | 566591 | 571159 | 4569 | reverse |
| Disease resistance protein RGA2 § | gene | 592710 | 594859 | 2150 | reverse |
| Putative disease resistance protein RGA1 § | gene | 604839 | 609329 | 4491 | reverse |
| Glycosyltransferase-like At3g57200 | gene | 644044 | 648953 | 4910 | forward |
| DNA-(apurinic or apyrimidinic site) lyase | gene | 670864 | 675667 | 4804 | forward |
| Glycosyl transferase | gene | 682384 | 687032 | 4649 | forward |
| U4/U6 small nuclear ribonucleoprotein PRP4-like protein | gene | 687890 | 692709 | 4820 | reverse |
| hypothetical protein TorRG33x02_272900 | gene | 700124 | 702379 | 2256 | reverse |
| hypothetical protein TorRG33x02_272900 | gene | 711204 | 713754 | 2551 | reverse |
| Probable inactive purple acid phosphatase 16 | gene | 828509 | 841426 | 12918 | forward |
| IQ motif, EF-hand binding site | gene | 841679 | 843725 | 2047 | forward |
| Glucan endo-1,3-beta-glucosidase, basic isoform § | gene | 845973 | 847034 | 1062 | reverse |
| Glucan endo-1,3-beta-glucosidase, basic isoform § | gene | 863658 | 867664 | 4007 | reverse |
| DYW domain containing protein | gene | 871587 | 872753 | 1167 | reverse |
| Glucan endo-1,3-beta-glucosidase, basic isoform § | gene | 880664 | 881873 | 1210 | reverse |
| Glucan endo-1,3-beta-glucosidase, basic isoform § | gene | 903147 | 904383 | 1237 | reverse |
| Glucan endo-1,3-beta-glucosidase, basic isoform § | gene | 915528 | 921253 | 5726 | reverse |
| RecA domain-containing protein | gene | 967798 | 970698 | 2901 | forward |
| Glucan endo-1,3-beta-glucosidase § | gene | 1001208 | 1002220 | 1013 | forward |
| F-box domain containing protein § | gene | 1011709 | 1013289 | 1581 | forward |
| Glucan endo-1,3-beta-glucosidase § | gene | 1032998 | 1034666 | 1669 | forward |
| LOB domain-containing protein 6 | gene | 1066960 | 1068347 | 1388 | forward |
| UDP-glucuronosyl/UDP-glucosyltransferase | gene | 1088695 | 1093385 | 4691 | reverse |

¶ - Not expressed
§ - Expressed

## DISCUSSION

This study utilized phenotyping and raw sequence data from Henning et al. (2017) and as such represents a limited evaluation of individuals for a genetic mapping and QTL study. Nevertheless, the sequencing depth for all individuals (48 individual genotypes per Illumina HiSeq 3000 sequencing lane) that were genotyped-by-sequencing was relatively high in comparison to previous GBS studies (Henning et al., 2016). In addition, the use of controlled environmental conditions in a glasshouse along with controlled inoculations resulted in consistent phenotypic values across replications. The combination of these two factors resulted in successful QTL analyses featuring an obvious QTL identified with little noise in non-QTL regions. Nevertheless, the most important contributions this study offered were twofold. First, our results demonstrated the superior genetic mapping made available by means of SNP identification using a PacBio reference genome over *de novo* genomes assembled by short-read sequencing. Secondly, the greater size of contigs from the PacBio genome (NG50 = 0.867 Mb) over that of genomes assembled from short-read sequencing (NG50 = 1.4 Kb) enabled clear identification of genes found nearby or within QTLs as opposed to partial or broken genes and empty space as seen in Henning et al (2017).

### Comparison Between Previous Map and Current map

Previous work by Henning et al. (2017) identified 10 linkage groups developed from 12098 initial markers that were filtered down to 2263 markers. In the development of this new genetic map, approximately 60K markers were filtered down to approximately 8k markers which were then used in Joinmap v5.0 to generate a map consisting of 2045 markers. In addition, the SNP set in Henning et al. (2017) was identified using the Teamaker genome assembly (Hill et al., 2017) while the SNP set used for this genetic map was identified using the Cascade PacBio sequenced and Falcon-assembled genome.

There are important differences between these two genomes as follows. First, the draft Cascade genome assembly (http://hopbase2.cgrb.oregonstate.edu/downloadDraftCascade.php) covers 4.24 Gb consisting of 11705 contigs, while the Teamaker genome only covers 2.8 Gb of the genome consisting of ~128K scaffolds with ~ 1.03 Gb of undetermined bases (Ns) connecting small contigs. Second, due to the small size of scaffolds in the Teamaker genome (NG50 = 1.4 kb), many genes are missing or broken between multiple scaffolds. In contrast, the Cascade genome consists of significantly larger contigs (NG50 = 866 kb) that are each long enough to contain hundreds of complete genes, gene families and their controlling elements.

Because SNPs were identified from a complete genome consisting of long contigs, the starting pool of SNP markers was significantly larger (60K vs 12K) and the number of high-quality markers that were available to develop linkage groups was also larger than Henning et al. (2017). These factors enabled the development of linkage maps clustering markers from the same contig together as well as had better saturation of markers along the whole linkage group. Only in the case of linkage group 1 was this not the case. This linkage group had one section, covering 14 cM to 45 cM, that averaged 3.4 cM between markers—significantly larger than the overall average distance of 0.32 cM between markers. All other linkage groups had consistent marker distribution along the maps.

The rational for developing genetic maps using ML for primary ordering of markers and use of REG with only two rounds of map estimation for relative marker order is backed by research from Hackett and Broadfoot (2003). This theoretical study looked at different mapping procedures where errors due to missing or misclassified markers were present. They identified ML as having the best statistics for ordering markers along a linkage group when compared to other methods. However, ML was significantly more sensitive than REG to missing genotypic data or misclassified genotypes when it came to estimating distances between markers. A combination of ML and REG mapping methods were used to overcome these difficulties and develop a genetic map with marker order being of greater import than distance between markers. Delimiting ML map distances to those relative to REG mapping with “2 rounds” of map estimation resulted in a map that presumably contains better marker order than REG and better estimation of genetic distances for each linkage group than ML. QTL mapping analyses are used to identify regions on linkage groups that are linked to expression of the trait under consideration. Genetic distances between QTLs are important for defining the probability of recombination and ultimately the success of obtaining favorable alleles at multiple QTLs. However, once a complete or near complete physical map with molecular markers is developed, genetic distances will be accurately predicted as will marker order.

The use of ML mapping to determine marker order, and use of REG for overall genetic distance of linkage groups, is one means for developing maps for QTL studies—particularly those involved in qualitatively controlled traits such as disease resistance. Use of combined ML and REG is particularly useful in situations where linkage groups contain 300 or more markers. In this case, one can run REG mapping with the “2 rounds” option selected for map development rather than opting for “3 rounds” as most of the computing time in JoinMap 5.0 occurs during the third round of map estimation. Use of “2 rounds” for map estimation in REG results in a linkage map having only a portion of the total number of markers available and in several cases resulted in adjacent markers on the same contig being separated into different positions on the linkage map. Nevertheless, a relatively accurate overall genetic distance for each linkage group can be obtained using “2 rounds” option. Then, delimiting the ML developed map by transforming distances between markers relative to the REG developed map obtained from “2 rounds” option, results in a linkage group with marker order that appeared to be a better match with the physical map along with reasonable distances between markers to determine overall genome coverage.

### Comparison Between Previous QTL Study and Current QTL study

Previous research (Henning et al., 2017) identified a single linkage group possessing three significant QTLs covering a range of 9 cM for PM resistance in this population. A limited analysis of genes found within and nearby QTLs did not reveal any R-genes known to be associated with disease resistance in other plant species although putative genes for chalcone synthase were identified. Part of the reason for missing R-genes or other known disease resistance genes in the previous study was a lack of a complete reference genome from which to identify SNP markers. As previously mentioned, the Teamaker & Shinsuwase genomes are missing up to an estimated 1 Gb of DNA assembly. It stands that these contigs observed in the present study were more than likely not included in the assembly and therefore would not be used for SNP identification. This would result in an imprecise identification of QTL along the linkage group and the potential for the identification of multiple QTLs spread across a wider region. In addition, we identified multiple putative genes associated with PM resistance—including R-genes. The significance of this finding is that no R-gene loci have been identified in hop to this date. Furthermore, the identification and upstream location of non-R-gene pathogenesis-related genes such as Glucan endo-1,3-beta-glucosidase and Peroxidase-27 provide hints for disease response targets that could potentially be used for constitutive-expression and quantitative disease resistance.

### Gene Identification

Three putative R-genes were identified on contig 002196F. A visual observation of RNAseq data (Padgitt-Cobb et al., 2019) obtained from USDA cultivar ‘Cascade’ (that was exposed to PM infection prior to collecting RNA samples) showed complex expression amongst these three with multiple variations of exons combined into putative isoform mRNA molecules with only two that appear to be complete transcripts (TCONS_00051515 and TCONS_00051522). The Augustus-predicted gene located at 002196F: 39443-41061 contains a full CC-NB-LRR complement exhibiting high similarity to <u>XP 024032005</u>, disease resistance protein RGA2-like [*Morus notabilis*]. The other putative R-gene located at 002196M: 46739-54401 have significant homology with <u>Q7XA39</u>: Putative disease resistance protein and <u>Q9LRR4</u>: Putative disease resistance RPP13-like protein. Other mRNAs expressed within 002196F; 39443 - 54401 regions appear to be putative partial R-genes. In addition to the two putative Augustus identified R-genes, contig 002196F contains a putative AP2-like ethylene-responsive transcription factor (AP2/ERF; <u>At2g41710</u>) expressed in Cascade cultivar tissues (Table 2). Fisher and Droge-Laser (2004) provide data showing that AP2/ERF mediate the expression of fungal pathogenesis-related genes in Tobacco. This transcription factor may provide a similar role in hop. Contig 002196F also contains a putative gene for polygalacturonase (<u>Q40312</u>) as well as a putative gene for Heat stress transcription factor B-1 (<u>Q22230.1</u>) (Table 2). The putative polygalacturonase gene was not expressed in any tissue while the heat stress transcription factor B-1 was expressed. Heat stress transcription factors have been shown to be active in response to abiotic stresses but have not generally been identified as responsive to biotic stresses such as pathogen infection (Guo et al., 2016). The potted plants of Cascade used for RNAseq in this study were not exposed to excessive heat, low water levels or salinity and it remains unknown why this transcription factor showed elevated levels of expression.

The seven putative R-genes found at 000559F:491369 – 609329 predicted by Augustus ranged in size from 2150 – 5980 nucleotides (Figure 4). As with the putative R-genes identified on contig 002196F, complex patterns of mRNA expression both within and among tissues were observed (Figure 4). Evaluation of the mRNA isoforms expressed from this putative R-gene locus show 19 different mRNAs (Table 4). Ten of these mRNAs share homology to other known R-genes. Only two of these transcripts (<u>TCONS_00020746</u> and <u>TCONS_00020755</u>) appear to fall within individual Augustus-predicted gene regions while possessing similar structure to one another with four exons. Both transcripts share homology with the same disease resistance protein RGA2 gene (<u>Q7XB09.1</u>). The other eight transcripts appear to have exons from multiple different Augustus-predicted genes and in some cases span ~89 kb’s of the 000559F contig.

**Figure 4.**
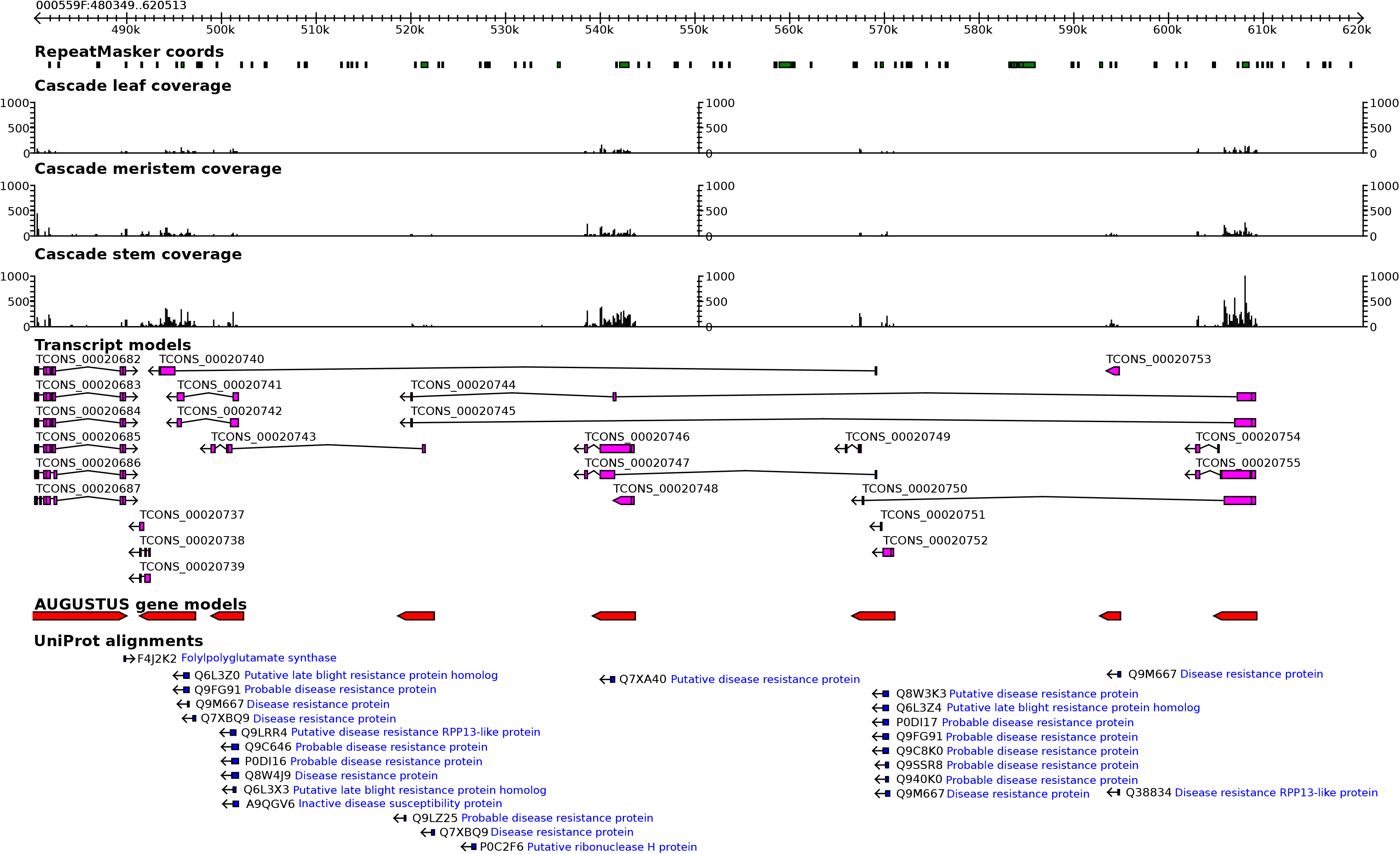
Extracted region of contig 000559F covering putative R-genes linked to expression of resistance to PM as shown in the Cascade draft genome on Hobpase.org (http://hopbase.cgrb.oregonstate.edu). Red color alignments represent Augustus-predicted genes while pink represents mRNA transcripts and blue represents Uniprot (https://www.uniprot.org) alignments. Quantitative expression levels (black color graphs) were all normalized between tissues.

**Table 4.** List of mRNA molecules expressed in putative R-gene locus. mRNA’s in red have significant homology to other known R-genes while the italicized, bold, red mRNA’s appear to be complete genes expressed within defined regions of the contig.

| mRNA Name | Start | Finish | Length | # Intervals |
| --- | --- | --- | --- | --- |
| TCONS_00020737 | 1 | 423 | 423 | 1 |
| TCONS_00020739 | 1 | 715 | 715 | 2 |
| TCONS_00020738 | 1 | 400 | 400 | 3 |
| TCONS_00020742 | 1 | 1203 | 1203 | 2 |
| TCONS_00020741 | 1 | 1204 | 1204 | 2 |
| TCONS_00020743 | 1 | 1163 | 1163 | 4 |
| TCONS_00020748 | 1 | 2231 | 2231 | 2 |
| ***TCONS_00020746*** | 1 | 3886 | 3886 | 4 |
| TCONS_00020749 | 1 | 236 | 236 | 3 |
| TCONS_00020740 | 1 | 1657 | 1657 | 3 |
| TCONS_00020747 | 1 | 1968 | 1968 | 4 |
| TCONS_00020751 | 1 | 233 | 233 | 1 |
| TCONS_00020752 | 1 | 1097 | 1097 | 2 |
| TCONS_00020753 | 1 | 1476 | 1476 | 1 |
| TCONS_00020754 | 1 | 376 | 376 | 2 |
| TCONS_00020744 | 1 | 2458 | 2458 | 4 |
| TCONS_00020745 | 1 | 2458 | 2458 | 3 |
| TCONS_00020750 | 1 | 3357 | 3357 | 3 |
| *TCONS_00020755* | 1 | 3960 | 3960 | 4 |

Eites and Dangl (2010) have demonstrated that in several cases two cis-acting R-genes are required for disease resistance. In our study, we observed two different putative “R-gene” loci that each contain what appear to be two fully functional R-genes. Our limited study cannot determine if the two adjacent putative R-genes on either contigs 002196F or 000559F act as suggested by Eites and Dangle (2010). Nevertheless, this work does identify potential molecular targets from which to design studies that could potentially answer such questions.

Unfortunately, the question concerning whether or not two R-genes are working in conjunction for resistance to PM is confounded by the presence of two potential R-genes (R4 and R6) loci in the same resistant parent genotype. The female parent of this cross, USDA Newport, appears to possess both R4 and R6 genes (Wolfenbarger et al. 2014) for resistance to PM. Later work by Wolfenbarger et al. (2016) suggests that R4 and R6 may be misclassified, the same gene, or so tightly linked that they segregate as a single group. Our observations on USDA Newport show two distinct regions that are separated by approximately 1 cM. Previous work by Wolfenbarger et al. (2016) could not distinguish between R4 and R6 genes in multiple genotypes possessing the R6 gene. It is possible that cultivar selection has centered on selecting lines that possess greatest fitness and these lines happen to contain both R4 & R6 genes. Test crosses such as the bi-parental cross used in our study could potentially expose both R-genes if they were not located directly adjacent to one another.

Several other pathogenesis-related genes were putatively identified using Augustus in addition to the putative R-genes found on contig 000559F (Table 3). Seven putative peroxidase-27 genes were identified along with seven putative Glucan endo-1,3-beta-glucosidase (GEBG) genes. None of the putative peroxidase genes were expressed while all of the putative GEBG genes were expressed or had portions of the Augustus delineated gene expressed (Figure 5). Again, complex expression patterns were observed for putative GEBG genes (Figure 5). In several cases (TCONS 00020766, TCONS 00020767, TCONS 00020768, TCONS 00020769) the transcripts are consisting of exons from multiple putative genes with TCONS 00020767 covering 52,763 nucleotides. All other transcripts appear to match up with Augustus-predicted genes. Differential quantitative expression of GEBG transcripts was observed with greatest expression of the transcript located at approximately 916K (Uniprot alignments: <u>Q02437</u>/<u>Q01412</u>) and some CEGB transcripts expressed in apical meristems and stems but not in leaves. Finally, we identified two Augustus-predicted putative F-box domain containing proteins (<u>PON47163.1</u>) with only one transcript located at nucleotide position 1011709 - 1013289 (<u>TCONS_00020702</u>) actually expressed. Previous studies (Van den Burg et al., 2008) have shown cell death and pathogen response for F-box proteins in Tobacco and Tomato. It is entirely possible this putative F-box protein is also involved in pathogenesis response to PM.

**Figure 5.**
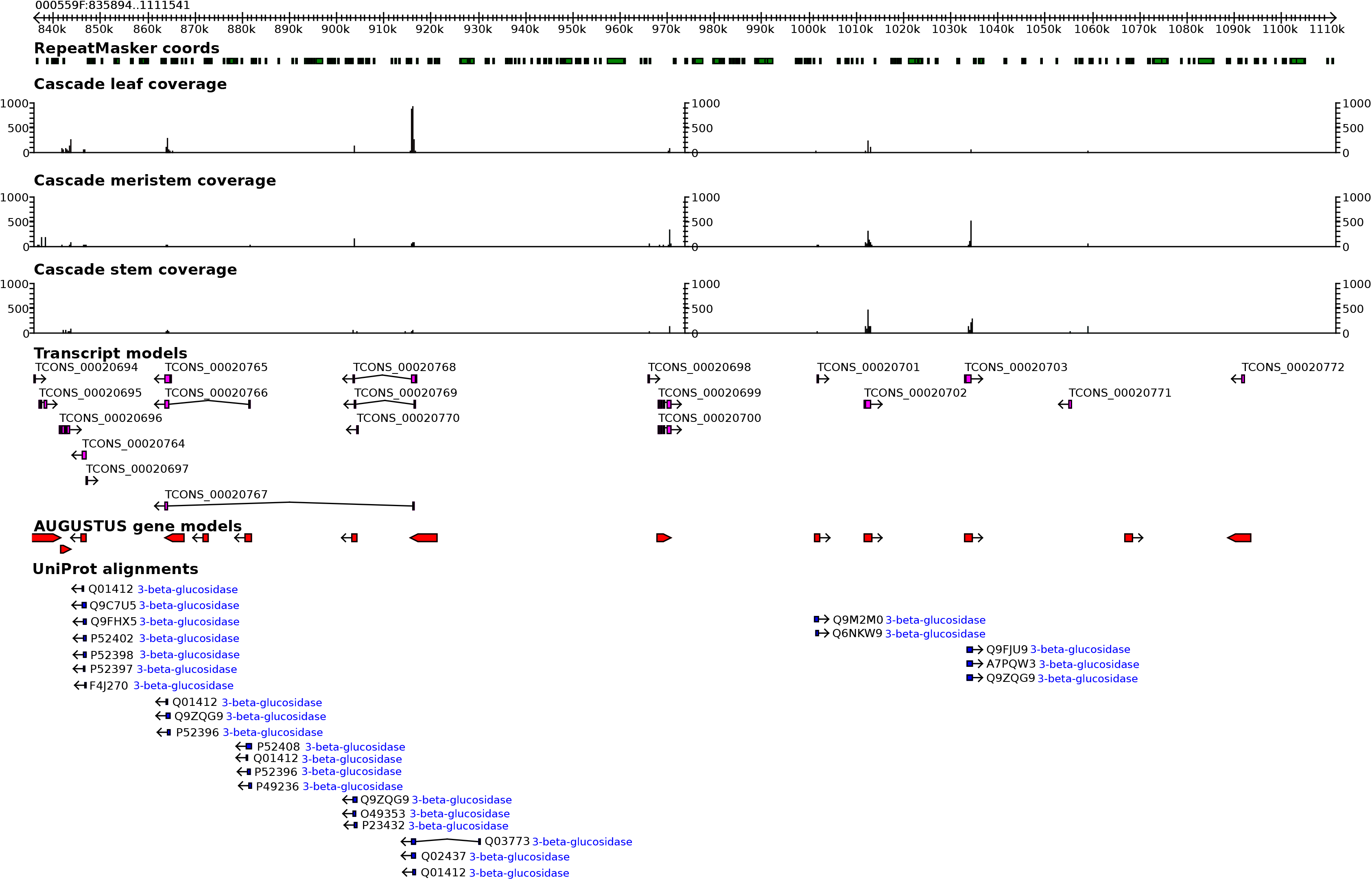
Extracted region of contig 000559F covering putative pathogen-response genes (Glucan endo-1,3-beta-glucosidase) linked to expression of resistance to PM. as shown in the Cascade draft genome on Hobpase.org (http://hopbase.cgrb.oregonstate.edu). Red color alignments represent Augustus-predicted genes while pink represents mRNA transcripts and blue represents Uniprot (https://www.uniprot.org/) alignments. Quantitative expression levels (black color graphs) were all normalized between tissues.

Our study identified a single QTL that covers two distinct, large contigs (002196F and 000559F) from the PacBio genome of Cascade and a bi-parental mapping population consisting of parents USDA Newport (PM resistant) and USDA 21110M (PM susceptible). Previous mapping studies (Henning et al., 2011 & 2017) were not able to clearly define genic regions, nor the genes themselves that were involved in pathogenesis response to PM. Our work was able to clearly define two regions of the hop genome—both located within 1 cM of each other—that actively respond to PM infection. In addition, for the first time in hop genomics, we can identify and define the genes and controlling factors that are involved in the expression of traits under genetic control along with their relative positions on linkage groups. This was not possible with short-read *de novo* assembled genomes due to the sheer number of small scaffolds and the resulting breakage of genes and gene families. Dependence upon genetic maps for determining and finding active genes involved in expression creates singular problems in that genetic maps rarely match 1:1 with physical maps. With significantly longer contigs from the PacBio genome, it is now possible to ascertain what genes are located next to significant molecular markers with more accurate gene prediction. In addition, annotation of the PacBio genome with RNAseq data facilitates an in-depth look at gene expression for traits of interest. With this in mind, it bears stating that gene expression—particularly that of putative R-genes—may be more complex than anticipated. In this study, we observed mRNA species that appear to be the result of “mix and match” between multiple regions of the putative R-locus to create putative R-genes, while other mRNA species are expressed from defined adjacent regions of the R-locus. Whether or not this observation represents artifacts of the transcript assembly or something analogous to human immune-responsive gene expression remains to be determined. More work is necessary to accurately define gene action for PM resistance— particularly work on additional R-genes in hop beyond R4 and R6.

## ACKNOWLEDGEMENTS

Funding for this study was provided by USDA-ARS CRIS project 2072-21000-051-00D. We wish to acknowledge Matthew Peterson and Chris Sullivan, CGRB-OSU, for their software and server support along with David Hendrix, Department of Biochemistry and Biophysics, OSU, for assistance with the final manuscript. Special thanks to Dr. David Gent (USDA-ARS-Corvallis, OR) Lab for inoculation and scoring disease resistance levels as reported in Henning et al. 2017. We would also like to thank Sierra Nevada Brewing for financial support and Pacific Biosciences for both financial and technical support for the development of the draft Falcon Cascade assembly.

